# Quantitative proteomics links the LRRC59 interactome to mRNA translation on the ER membrane

**DOI:** 10.1101/2020.03.04.975474

**Authors:** Molly M. Hannigan, Alyson M. Hoffman, J. Will Thompson, Tianli Zheng, Christopher V. Nicchitta

## Abstract

Protein synthesis on the endoplasmic reticulum (ER) requires the dynamic coordination of resident membrane proteins and cytoplasmic translation factors. While ER membrane proteins functioning in ribosome association, mRNA anchoring, and protein translocation, have been identified, little is known regarding the higher order organization of ER-localized translation. Here we utilized proximity proteomics to identify neighboring protein networks for the ribosome interactors SEC61β, RPN1, SEC62, and LRRC59. Whereas the SEC61β and RPN1 BioID reporters revealed translocon-associated networks, the SEC62 and LRRC59 reporters identified divergent interactome networks of previously unexplored functions. Notably, the SEC62 interactome is enriched in redox-linked proteins and ER luminal chaperones, whereas the LRRC59 interactome is enriched in SRP pathway components, translation factors, and ER-localized RNA-binding proteins. Analysis of the LRRC59 interactome by native immunoprecipitation identified similar protein and functional enrichments. Combined, these data reveal a functional domain organization for the ER and suggest a key role for LRRC59 in the organization of mRNA translation on the ER.

**Summary:** Hannigan et al. characterize the protein interactomes of four ER ribosome-binding proteins, providing evidence that ER-bound ribosomes reside in distinct molecular environments. Their data link SEC62 to ER redox regulation and chaperone trafficking, and suggest a role for LRRC59 in SRP-coupled protein synthesis.

## Introduction

RNA localization and accompanying local translation serve critical roles in the spatiotemporal regulation of post-transcriptional gene expression. Reflecting the importance of such regulation, localized mRNA translation requires the coordinate localization of numerous proteins, including aminoacyl-tRNA synthetases, translation factors, RNA-binding proteins (RBPs), molecular chaperones, enzymes/scaffolding proteins which act to modify the nascent polypeptide chain, as well as *cis*-encoded mRNA localization and trafficking information (Bellon et al., 2017; Debard et al., 2017; Gu et al., 2004; Gunkel et al., 1998; Huttelmaier et al., 2005; Koppers et al., 2019; Micklem et al., 2000; Paquin et al., 2007; Smibert et al., 1999; Tiruchinapalli et al., 2003; Vidaki et al., 2017; Willett et al., 2011; Yasuda et al., 2013; Zhang et al., 2017). At the endoplasmic reticulum (ER), the primary site for secretory and membrane protein synthesis, mRNA translation becomes even more complex, requiring additional protein factors including proteins that facilitate ribosome association with the ER membrane, which includes the translocon itself, and newly discovered non-canonical integral membrane RNA-binding proteins (Beckmann et al., 2001; Berkovits and Mayr, 2015; Cui et al., 2012; Gorlich et al., 1992; Hsu et al., 2018; Jagannathan et al., 2014; Jan et al., 2014; Johnson and van Waes, 1999; Rapoport, 2007; Reid and Nicchitta, 2012; Reid and Nicchitta, 2015a; Simsek et al., 2017; Stephens et al., 2005; Voigt et al., 2017; Walter, 1981a; Walter, 1981b).

An additional level of complexity to the organization of ER-localized protein synthesis appears when considering the multiple lines of evidence that support a transcriptome-wide role for the ER in proteome expression (Chartron et al., 2016; Cui et al., 2012; Diehn et al., 2006; Diehn et al., 2000; Hoffman et al., 2019; Jan et al., 2014; Lerner et al., 2003; Mueckler and Pitot, 1981; Mueckler and Pitot, 1982; Reid and Nicchitta, 2012; Reid and Nicchitta, 2015a; Voigt et al., 2017). Notably, investigations of ER-localized mRNA composition in human cells, tissues, yeast, and fly revealed that all transcripts, not just those encoding secretory and membrane proteins, are translated on the ER (Chartron et al., 2016; Chen et al., 2011; Cui et al., 2012; Diehn et al., 2000; Jan et al., 2014; Kopczynski et al., 1998; Lerner et al., 2003; Mueckler and Pitot, 1981; Mueckler and Pitot, 1982; Reid and Nicchitta, 2012; Reid and Nicchitta, 2015a; Voigt et al., 2017). While landmark biochemical and structural studies have advanced our understanding of how secretory/membrane protein synthesis is coupled to protein translocation, it remains unclear how translation on the ER is compartmentalized to accommodate the coincident translation of both cytosolic and secretory/membrane protein-encoding mRNAs. One model proposes that an mRNA-wide role for the ER in proteome expression is achieved by translocon-independent modes of ribosome association with the ER membrane (Harada et al., 2009; Kreibich et al., 1978a; Levy et al., 2001; Muller and Blobel, 1984; Reid and Nicchitta, 2015a; Savitz and Meyer, 1993; Tazawa et al., 1991). In this view, the SEC61 translocon serves a canonical role in secretory/membrane protein biogenesis by recruiting ribosomes engaged in the translation of this mRNA cohort, while other candidate ribosome interactors (e.g., p180, p34/LRRC59, SEC62) function as non-translocon ribosome binding sites. Ribosomes bound at these non-translocon sites may engage in the translation of both cytosolic and secretory/membrane protein-encoding transcripts. In the case of secretory/membrane polypeptides undergoing early elongation on non-translocon-associated ribosomes, we postulate a process where signal sequence-bearing nascent chains access translocons via lateral diffusion (Chartron et al., 2016; Jan et al., 2014; Jan et al., 2015; Reid and Nicchitta, 2015a; Reid and Nicchitta, 2015b). A primary prediction of this model is that different ribosome interacting proteins would reside in distinct membrane protein environments, perhaps reflecting the degree to which their bound ribosomes are dedicated to secretory/membrane protein synthesis. With understanding of the structural organization and regulation of ER-associated translation being largely derived from the classical canine pancreas rough microsome system, a largely unexplored question in the field thus concerns the cellular components and mechanisms that support the diversity of ER-localized translation in the cell.

Using a BioID proximity-labeling approach to examine this model, we recently reported that SEC61β, a translocon subunit, and the candidate ribosome-binding protein LRRC59 interact with populations of ribosomes engaged in the translation of divergent cohorts of mRNAs (Hoffman et al., 2019). In this communication, we extend these studies by investigating the ER protein interactomes of the four previously engineered BioID reporters (SEC61β, RPN1, SEC62, and LRRC59) (Hoffman et al., 2019). In time course labeling studies, we observed that for each reporter, proximal interactome labeling intensified but only modestly diversified as a function of labeling time, a finding consistent with a functional domain organization of the ER. Unexpectedly, our data revealed that the previously reported ribosome receptor SEC62 interacts with unique and unexpected protein networks, including those with roles in cell proliferation, signaling pathways, redox homeostasis, and cytoplasmic displaced ER luminal chaperones. In contrast, LRRC59 displays a highly SRP pathway-, translation-, and RNA-binding protein-enriched interactome. Both proximity proteomics and native immunoprecipitation studies found LRRC59 to interact almost exclusively with SRP machinery, non-canonical ER-RBPs, and translation initiation factors, suggesting a previously unappreciated role for LRRC59 in the organization and/or regulation of secretory/membrane protein synthesis on the ER.

## Results

### Evidence for domain organization of ER membrane protein interactomes

In a recent study, we examined the spatial organization of mRNA translation on the endoplasmic reticulum via proximity proteomics, where BioID reporters of translocon-associated (SEC61β, RPN1) and candidate (SEC62, LRRC59) ribosome interacting proteins were used to biotin label proximal ribosomes *in vivo*. Together with RNA-seq analysis of mRNAs isolated from the biotin-tagged ribosome populations (Hoffman et al., 2019), these studies revealed that translation on the ER membrane is heterogeneous and that ER-bound ribosomes display local environment-specific enrichments in their associated mRNAs. The mechanism(s) responsible for this regional organization of translation, however, is unknown. Here, we used proximity proteomics and the previously utilized BioID reporters to test the hypothesis that ribosome-binding proteins reside in distinct interactome networks or functional domains, as a potential mechanism to support higher order organization of mRNA translation on the ER.

In the experiments presented below, BioID reporters of known ribosome interacting proteins were used to map proximal ER membrane protein interactomes at previously identified mRNA translation sites (Figure 1A) (Hoffman et al., 2019). BioID proximity labeling experiments are typically conducted over many hours (Roux et al., 2012; Sears et al., 2019; Varnaite and MacNeill, 2016) (e.g. 16-24 hours), a reflection of the slow release kinetics of the reactive biotin-AMP catalytic intermediate from the BirA* active site (Kwon and Beckett, 2000). In context of this study, we considered that such extended labeling times, coupled with reporter diffusion in the ER membrane, would confound identification of proximal-interacting vs. random-interacting proteins. In line with this consideration, we expected that for each reporter, the composition of biotin-tagged proteins would diversify as a function of labeling time (Rees et al., 2015). Though it has been previously demonstrated that neighboring interactomes can be distinguished from random interactors by their higher relative labeling over non-specific controls, we first examined timecourses and patterns of biotin labeling for the BioID reporters noted above (Kim et al., 2014; Rees et al., 2015; Roux et al., 2012). The results of these experiments are shown in Figure 1B. Depicted are streptavidin blots of the cytosol (**C**) and membrane (**M**) protein fractions from the four BioID reporter cell lines, sampled over a labeling time course of 0–6 hours. Two observations are highlighted here. One, although the BirA domains are cytosolically disposed, biotin-tagging is strongly enriched for membrane vs. cytosolic proteins. Two, the major membrane protein biotin labeling patterns intensify but did not substantially diversify over the labeling time course (Figure 1B). Densitometric analysis of the biotin labeling patterns revealed by SDS-PAGE are depicted in Figure 1B, right panels, where it can be further appreciated that the overall labeling patterns were relatively constant over labeling time. These data suggest that the BioID interactomes of the tested reporters include largely stable membrane protein assemblies, rather than the randomizing interactomes expected of diffusion-based interactions (Goyette and Gaus, 2017; Kusumi et al., 2012; Kusumi et al., 2011; Singer and Nicolson, 1972). The data presented above (Figure 1B) are consistent with a model where the local environments of the BioID reporters are constrained. Such spatial restriction may reflect an organization of the ER via functional interactome networks, similar to the well documented observations of plasma membrane domain organization (Goyette and Gaus, 2017; Kusumi et al., 2012; Kusumi et al., 2011). We also considered that the distinctive labeling patterns of the different reporters could be influenced by ER dynamics and/or distribution biases of the reporters (e.g. tubules vs. lamellar regions). To examine these scenarios, we performed BirA* labeling time course experiments *in vitro*, using canine pancreas rough microsomes (RM) which lack the native topology and dynamics of the ER, and a soluble, recombinant BirA* (Figure 1C). Using this experimental system, the reactive biotin-AMP intermediate was delivered in *trans* and accessible to the microsome surface by solution diffusion. The results of these experiments demonstrate that when accessible to RM proteins in *trans*, biotin labeling is pervasive, with RM proteins being broadly labeled and labeling intensities increasing as a function of labeling time (Figure 1C, upper panel; protein loading control depicted in Figure 1C, lower panel). Combined, the distinct and temporally stable proximity labeling patterns identified for each BioID reporter cell line suggest that the BirA-chimeras reside in distinct protein interactome domains of the ER.

**Figure 1.**
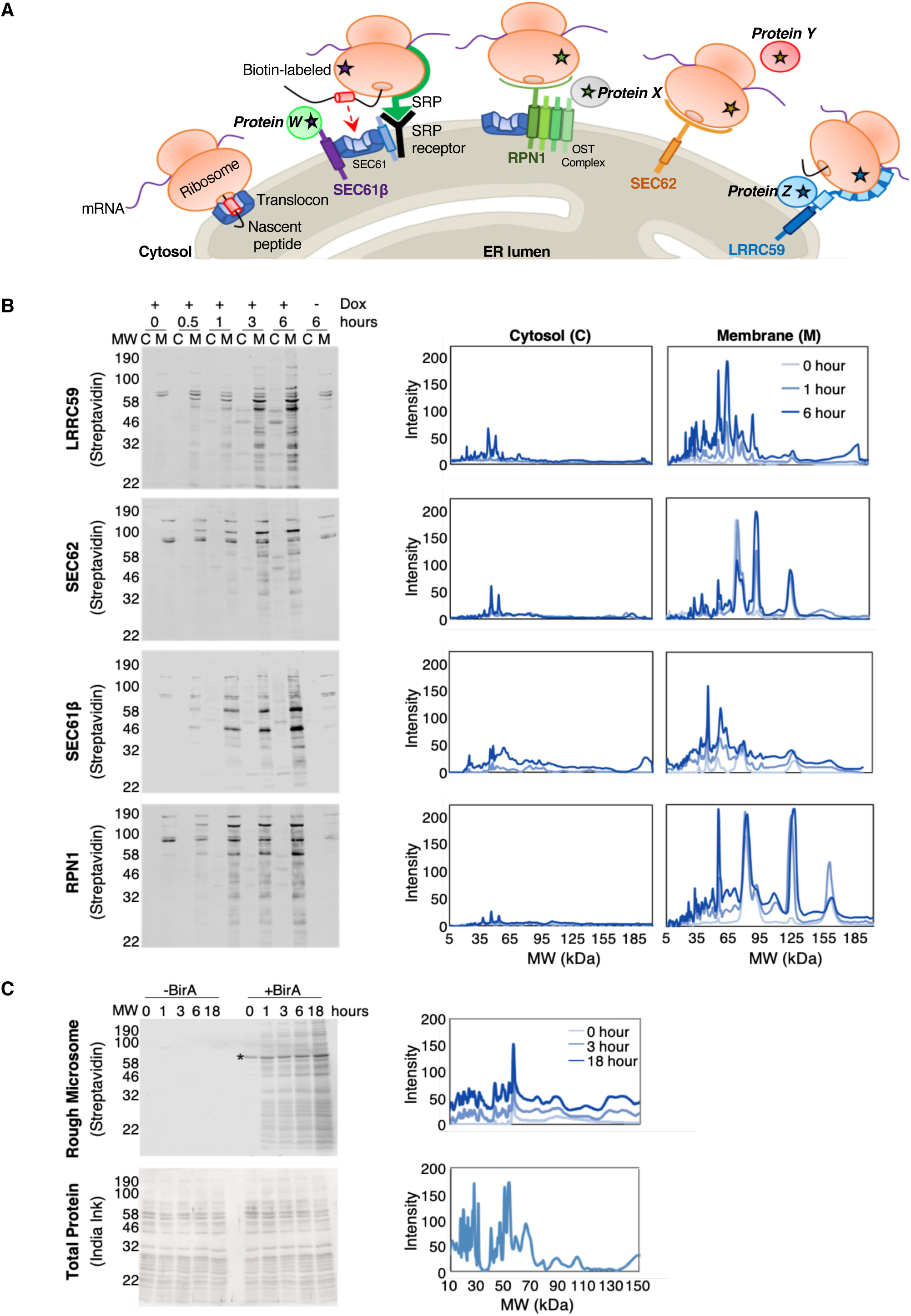
Identification of ER membrane protein interactomes by proximity proteomics. **(A)** Schematic of known (SEC61 translocon, OST complex), and candidate (SEC62, LRRC59) ER-ribosome receptors. SEC61β (purple), a subunit of the SEC61 translocon, RPN1 (green), a subunit of the OST complex, SEC62 (orange), and LRRC59 (blue) are expressed as BioID chimeras, labeling interacting and near-neighbor proteins (indicated by starred ribosomes and proteins W, X, Y, and Z). **(B)** Left panel: Streptavidin blots examining the subcellular distribution of biotin-labeled proteins within HEK293 cells expressing either the LRRC59-, SEC62-, SEC61β-, or RPN1-BirA reporter constructs. Biotin labeling (doxycycline (dox)-inducible expression of reporters) was performed over a time course spanning 0-6 hours and cytosol (C) and membrane (M) extracts prepared by detergent fractionation. Right panel: Densitometric quantifications of biotin labeling intensities for cytosolic and membrane fractions. **(C)** Canine pancreas rough microsomes with (+BirA) or without (-BirA) the addition of BirA* in *trans*. Biotin labeling of proteins was conducted over 0-18 hours (top, left). Biotin labeling intensities were quantified using densitometric analyses (top, right). As a loading control, total protein lysate was analyzed by India ink staining (bottom, left) and quantified by densitometric analysis (bottom, right).

### Investigation of local interactomes via TMT quantitative mass spectrometry

To enable quantitative measurements of the protein interactomes schematically illustrated in Figure 1, an isobaric-tagging mass spectrometry analytical approach was used (TMT: tandem mass tagging) (Figure 2A). Isobaric labeling methods provide multiplexing and, in this case, quantitative analysis of biological replicates, enhancing the reproducibility and accuracy of datasets. Two oligomeric protein complexes known to reside at sites of translation on the ER, the SEC61 translocon and the oligosaccharyltransferase (OST) complex, were used as spatial reference points with the expectation that they would label their associated subunits (Figures 1A, 2A). Specifically, for the SEC61 translocon, a BioID reporter of its subunit, SEC61β, was used to map the interactome of this well-studied complex (Becker et al., 2009; Beckmann et al., 2001; Dejgaard et al., 2010; Pfeffer et al., 2015; Voorhees et al., 2014). Similarly, ribophorin I (RPN1), a subunit of the OST complex that is transiently recruited to the SEC61 translocon during nascent glycoprotein translocation, served as a parallel proximity labeling control for the local environments of ER translation sites (Figures 1A, 2A) (Kelleher et al., 1992; Kreibich et al., 1978a; Nilsson et al., 2003; Wild et al., 2018). To expand our analysis to less studied ER environments, we examined LRRC59 as it has been previously reported to reside proximal to ER-bound ribosomes *in vivo* (Hoffman et al., 2019) and to function in ribosome binding*in vitro* (Ichimura et al., 1993; Tazawa et al., 1991) (Figure 1A). We also investigated a second candidate ribosome-binding protein, SEC62, which has been demonstrated to bind ribosomes in vitro and to be in the vicinity of bound ribosomes in permeabilized cell models (Hoffman et al., 2019; Lang et al., 2012; Muller et al., 2010). While both LRRC59 and SEC62 have been shown to interact with ribosomes, their native protein interactomes are largely unstudied.

**Figure 2.**
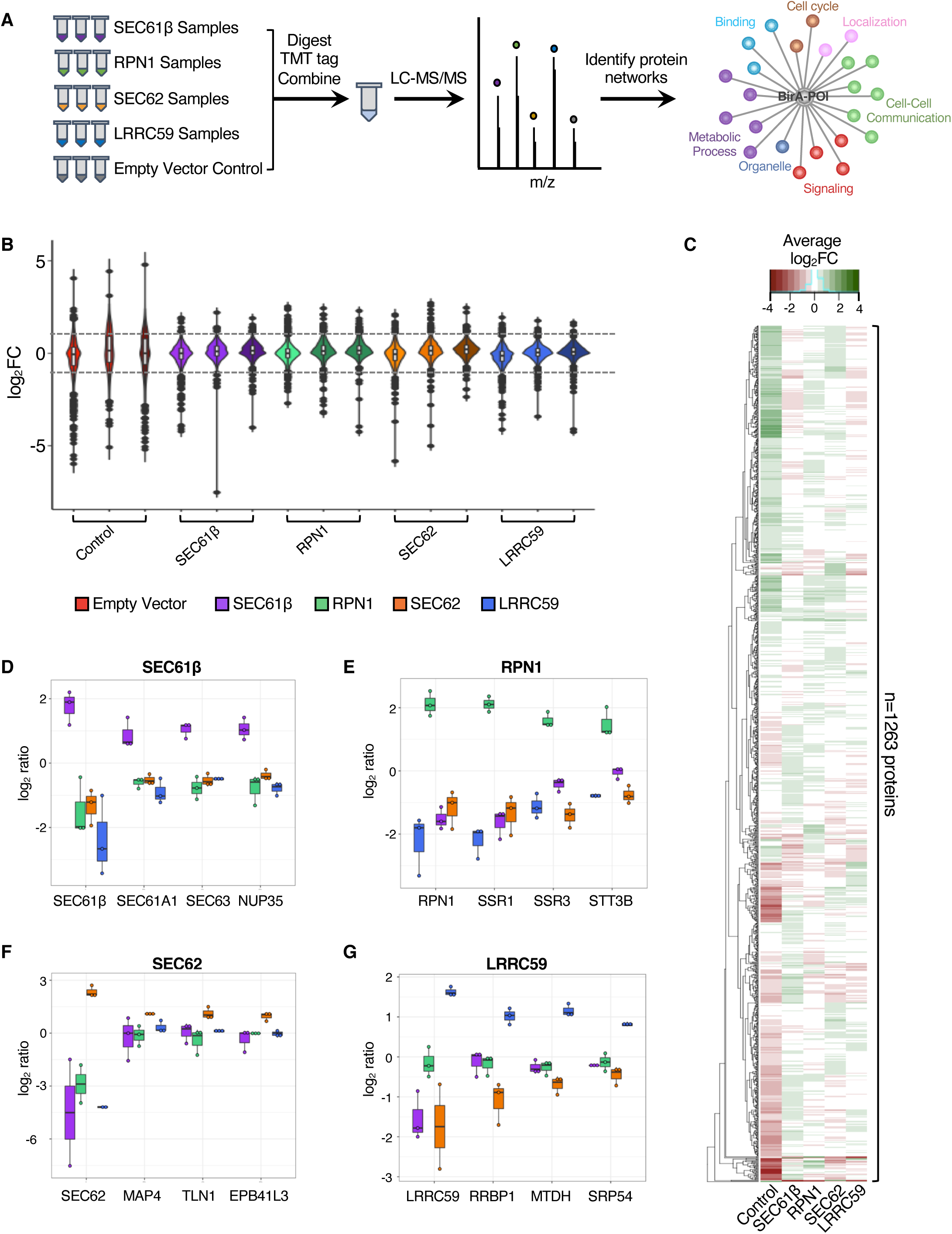
Proximity proteomics reveals unique interactomes for each of the four tested baits. **(A)** Schematic of the experimental approach. BirA-reporters for known (SEC61β (purple), RPN1 (green)) and candidate (SEC62 (orange), LRRC59 (blue)) ER-resident ribosome interacting proteins were expressed with biotin labeling (3 hours) conducted in biological triplicate. An empty vector negative control (red) was included. Samples were digested, tandem mass tag (TMT) labeled, and combined for liquid chromatography-tandem mass spectrometry (LC-MS/MS) analysis. Enrichment analyses of biotin-labeled proteins will reveal protein-protein interactions and/or functional networks for each of the five baits. **(B)** Violin plots of the protein abundance distributions for all biotin-labeled proteins (n=1,263) for each bait. **(C)** Clustered heatmap showing the average log_2_FC (across biological replicates) for each of 1,263 identified proteins per bait (green represents enriched protein abundance; red indicates decreased protein abundance). Boxplots showing the enrichment of highly enriched proteins (prey) labeled in the **(D)** SEC61β, **(E)** RPN1, **(F)** SEC62, and **(G)** LRRC59 BioID reporter studies. Each dot represents the log_2_ FC value per biological replicate.

We established inducible Flp-In™ T-Rex™ HEK293 cell lines for each of the BioID reporters and included an empty vector negative control Flp-In™ T-Rex™ HEK293 cell line for background characterization. By the rationale detailed above, cell lines were biotin-labeled for three hours to allow for significant labeling of intracellular membrane proteins (Figure 1B), affinity isolated from cell extracts, digested with trypsin, derivatized with isobaric mass tag reagents, combined, and analyzed by LC-MS/MS for identification of protein networks (Figure 2A). To enable the analysis of three biological replicates for each of the four cell lines, in addition to six study pool QC replicates, two TMT 10-plex reagent sets were utilized. Biological groups were divided between the TMT sets to avoid between-set bias, and the SPQC replicates were used to normalize between TMT sets.

### Identification of ER membrane protein interactomes

Quantification and identification of TMT-labeled peptides for each of the different BioID reporters were performed with Protein Discoverer 2.3 and Scaffold Q+ software. TMT signals were normalized to the total intensity within each channel, peptides derived each protein summed to represent the protein abundance, and relative protein abundance was calculated as a log_2_ fold change (FC) relative to the mean of the SPQC reference channels, which represents the biological average of all samples in the experiment. In total, 1,263 proteins were identified across the entire sample set, with the majority of proteins showing modest to no reporter-specific enrichment (Figure 2B, **Supplemental File S1**). Violin plots in Figure 2B highlight the technical reproducibility of the approach. Despite SEC61β, RPN1, SEC62 and LRRC59 sharing similar overall log_2_FC distribution patterns (Figure 2B), examination of the magnitude of biotin labeling at the protein level revealed that each reporter is associated with a unique set of prominent near-neighbor interactors (log_2_FC > 1, dashed line), as summarized in the heatmap profile (Figure 2C), and individual reporter representations (Figure 2D-G). As depicted, the SEC61β reporter labeled other members of the SEC61 translocon, as well as a nuclear pore complex protein (Figure 2D); the RPN1 reporter labeled subunits of the OST complex and other glycoproteins (Figure 2E); the SEC62 interactome includes an array of proteins involved in redox regulation, cytoskeleton architecture, and the cell cycle (Figure 2F); and the LRRC59 interactome included ribosome-binding proteins, RNA-binding proteins, and SRP pathway components (Figure 2G). Importantly, all of the bait proteins significantly labeled themselves, providing a quantitative index of relative proximity (Figure 2D-G). Since identification and quantification are not decoupled in isobaric tagging experiments (e.g. the identification and quantification come from the same spectrum, which is a mixture of all samples), we also performed BirA-reporter proteomic studies using label-free shotgun proteomics (not multiplexed). Although this approach did not have the proteome coverage of the TMT-tagging approach, we were able to independently verify the high-confidence interactors for each reporter. Specifically, we identified SEC61 subunits, members of the OST complex, factors related to redox homeostasis and the cytoskeleton, and an enrichment of SRP machinery, translation factors and RBPs in the SEC61β, RPN1, SEC62, and LRRC59 interactomes, respectively, using this approach (Supplemental Figure 1C, **Supplemental File S3**). Combined, these data indicate that ER proteins can reside in discrete protein interactomes, which is consistent with a model where cohorts of functionally-related or interacting proteins comprise stable membrane domain interactomes, as previously reported for other membrane systems (de Brito and Scorrano, 2010; English and Voeltz, 2013; Helle et al., 2013; Hung et al., 2017)

### Characterization of SEC61β and RPN1 interactomes using proximity proteomics

To further characterize the protein interactomes of the reporter baits, we combined statistical prioritization, 2D clustering, and principal components analysis. This integrative approach bypasses the somewhat arbitrary requirement of filtering against a specific fold-change value, and instead uses protein co-expression patterns to identify interaction networks, thereby correct for variability in protein abundance across each of our reporter cell lines. This analysis identified 145, 13, 50, and 25 high-confidence protein interactors of SEC61β, RPN1, SEC62, and LRRC59, respectively (**Supplemental File S2**). Since the interactomes of SEC61β and RPN1 are at least in part characterized, we first examined the protein networks of these two baits. In this analysis, SEC61β had the highest number of high-confidence interactors (n=145 proteins) (Figure 3A), making it the largest interactome captured by our study. Despite its large size, gene ontology (GO) analysis demonstrated that nearly all of SEC61β protein partners (either direct or proximal) are membrane proteins and/or have functions related to protein transport (Figure 3B), which aligns with the known functions of SEC61 in ER targeting, membrane insertion, and translocation of newly synthesized polypeptides (Lang et al., 2017). Moreover, almost half (44%) of the identified protein interactors are annotated to physically interact with one another, suggesting that the SEC61β interactome is not only enriched for membrane/secretory proteins but that these high-confidence interactors comprise large protein-protein complexes/networks (Figure 3C). Notably, our proteomics and protein-protein interaction (PPI) analyses revealed that SEC61β interacts with SEC61*α* (SEC61A1) and SEC63 (Figures 2D, 3C), which is consistent with previous reports (Becker et al., 2009; Gorlich et al., 1992; Hartmann et al., 1994; Lang et al., 2012) and further validates that the putative interactors identified by our approach are likely *bona fide* targets.

**Figure 3.**
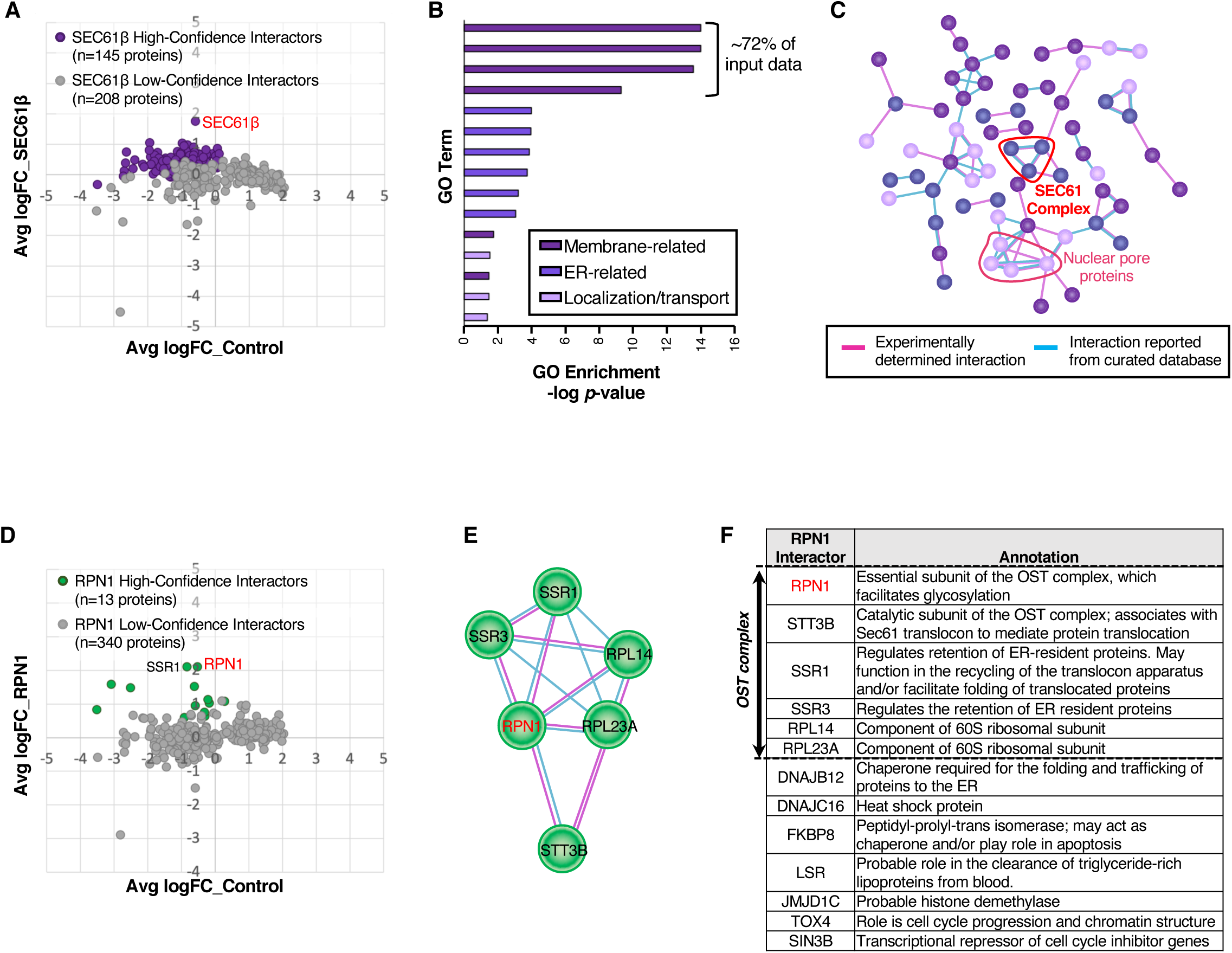
Characterization of protein networks for known ribosome interactors, SEC61β and RPN1. **(A)** Comparison of protein abundance for the 353 identified putative interactors in SEC61β-BirA and empty vector control HEK293 cells. Purple dots represent enriched, high-confidence interactors. Gray dots represent proteins that are less likely *bona fide* interactors of SEC61β-BirA. **(B)** Enriched Gene Ontology (GO) terms associated with high-confidence SEC61β-BirA interactors. Dark purple, purple, and light purple bars represent membrane-, endoplasmic reticulum (ER)-, and protein transport-related GO enriched terms, respectively. **(C)** Protein-protein interactions (PPI) among high-confidence SEC61β-BirA interactors, based on STRING annotations. Pink and cyan edges indicate experimentally determined and curated interactions, respectively. **(D)** Comparison of protein abundance for the identified 353 putative interactors in RPN1-BirA and empty vector control HEK293 cells. Green dots represent enriched, high-confidence proteins that interact with RPN1-BirA. Gray dots represent proteins that are less likely *bona fide* interactors of RPN1-BirA. **(E)** PPI network among high-confidence RPN1-BirA interactors, based on STRING annotations. **(F)** Functional comparison of all 13 high-confidence RPN1-BirA interactors, based on STRING annotations.

In contrast to the large number of proteins identified as SEC61β interactors, examination of the RPN1 interactome yielded the smallest number of interactors (n=13 proteins) (Figure 3D). Despite its small size, about one-third of the RPN1 interactome comprises members of the OST complex (Figure 3E-F), including STT3B, and the *α* and β subunits of the TRAP complex (SSR1, SSR3), as expected (Nilsson et al., 2003; Pfeffer et al., 2014). Additionally, our analysis revealed RPN1 to interact with 60S ribosomal proteins (RPL14, RPL23A), supporting a role for RPN1 in ribosome association (Braunger et al., 2018). Collectively, our characterizations of the SEC61β and RPN1 interactomes parallel high-resolution structural analyses of the SEC61 translocon, which place the OST and TRAP complexes in close physical proximity to the SEC61 oligomer (Nilsson et al., 2003; Pfeffer et al., 2014).

### Functional diversity across the BioID-SEC62 interactome

Following the statistical methodology described above, we interrogated the SEC62 interactome. As mentioned earlier, SEC62 has been demonstrated to interact with ribosomes and to facilitate mRNA translation and protein translocation on the ER (Lang et al., 2012; Muller et al., 2010); however, a comprehensive understanding of the SEC62 interactome in mammalian cells has not been previously reported. As assessed by BioID proteomics, the SEC62 interactome of HEK293 cells is comprised of a large cohort of proteins (n=50) (Figure 4A). Consistent with our previous study (Hoffman et al., 2019), we did not identify significant interactions of the SEC62 reporter with ribosomes, indicating that SEC62 may participate in ER translation independent of ribosome binding, as postulated for the canine pancreas rough microsome system (Jadhav et al., 2015).

**Figure 4.**
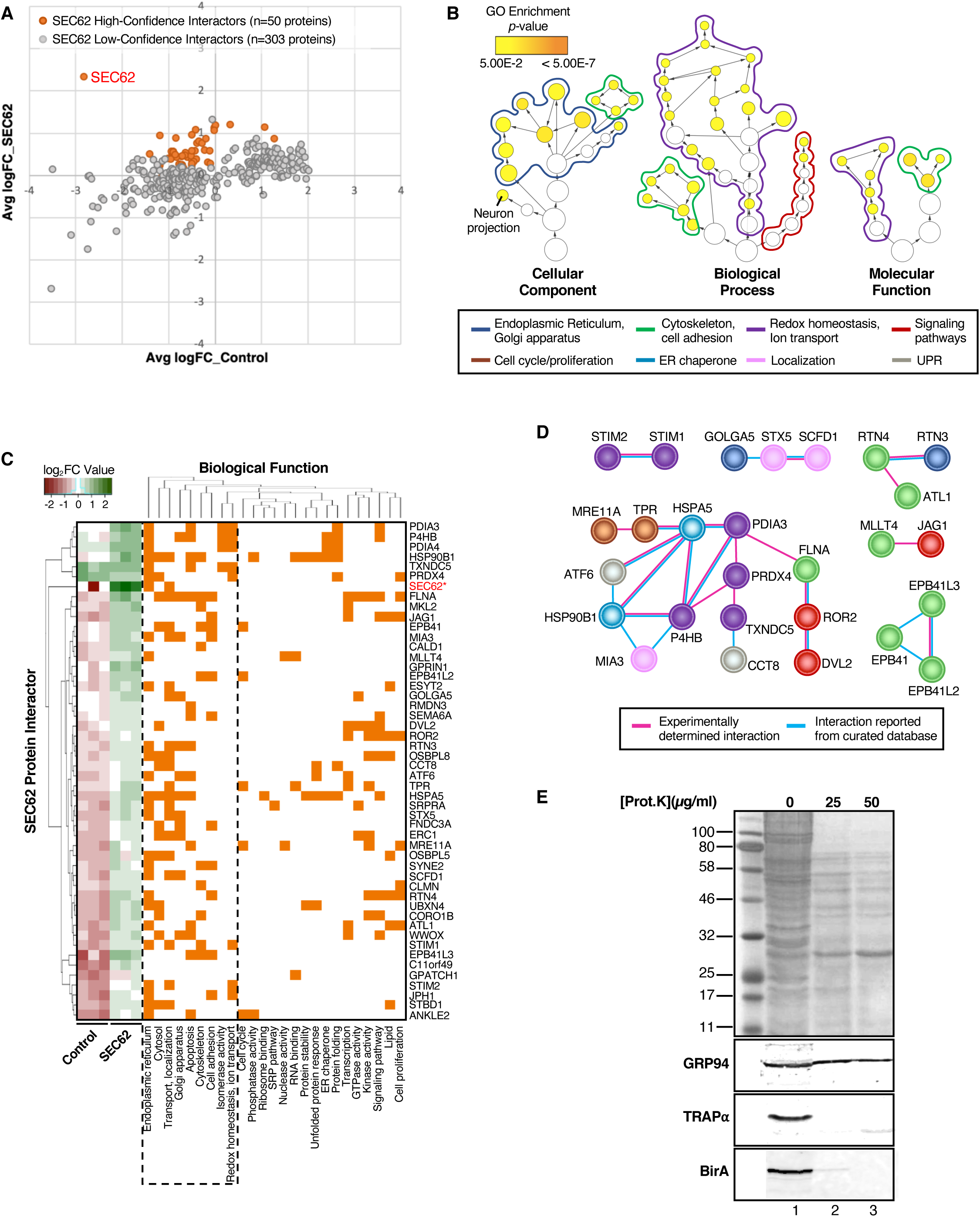
BioID-SEC62 labels functionally diverse proteins. **(A)** Comparison of protein abundance for the 353 identified putative interactors in SEC62-BirA and empty vector control HEK293 cells. Orange dots represent enriched, high-confidence proteins that interact with SEC62-BirA. Gray dots represent proteins that are less likely *bona fide* interactors of SEC62-BirA. **(B)** Hierarchical view of relationships for GO terms associated with SEC62 high-confidence protein interactors. GO term circles are outlined to match the colors assigned to each enriched GO category, as indicated beneath the panel. Circle sizes represent the number of genes in each enriched term, whereas circle color indicates the GO enrichment *p*-value. **(C)** Clustering of SEC62 high-confidence interactors based on co-occurrence of functional annotations. The left-most heatmap represents protein abundance values across biological replicates in control and SEC62-BirA HEK293 cells. **(D)** Protein-protein interactions (PPI) among high-confidence SEC62-BirA interactors, based on STRING annotations. Proteins are color-coded to match their functional assignment, as indicated above the panel. **(E)** Topology analysis of SEC62-BirA reporter line. SEC62-BirA cultures were chilled on ice, permeabilized with a digitonin-supplemented cytosol buffer, and subjected to digestion with the indicated concentrations of Proteinase K for 30 min on ice. Cells were subsequently lysed and total protein was resolved via SDS-PAGE (top panel). Following transfer, membranes were probed for GRP94 (ER-luminal protein), TRAP*α* (ER-resident protein with cytosolically-disposed antibody epitope), and BirA (BioID-SEC62 reporter). Lanes 1, 2, and 3 represent digestions with 0, 25, and 50 μg/ml proteinase K, respectively.

Alternatively, the BioID reporter construct may occlude ribosome binding activity present in the native protein. To examine the functional significance of the BioID-SEC62 interactome, we performed GO analysis and database mining on the 50 identified proteins. This analysis revealed that SEC62 interacts with a wide range of proteins involved in biologically diverse functions, including roles in cell cycle and proliferation, cytoskeleton architecture, protein localization, signaling pathways, ER chaperones, and redox homeostasis (Figure 4B-C). Since SEC62-interacting proteins have overlapping cellular functions (Figure 4C), we next asked if these proteins physically interact with one another to form protein complexes that may provide mechanistic insights into the biological functions of the SEC62 interactome. Protein-protein interaction analysis revealed that 54% of the SEC62 interactome physically interact with one another (Figure 4D). Using literature-based searches, database mining, and informatic approaches, we assigned a primary function to each protein, as indicated by the color legend in Figure 4B. In this depiction, the edges connecting interacting proteins were color coded to distinguish experimentally determined interactions from those reported/curated in databases (cyan), as annotated by the STRING database (Szklarczyk et al., 2019; Szklarczyk et al., 2017). Similar to the GO analysis, interrogation of PPI networks demonstrated heterogeneity in the functional assignment of interacting proteins. Notably, SEC62 was not reported in any of the six PPI networks, which further emphasizes the current lack of knowledge regarding SEC62 interactions in cells.

A particularly striking finding in the SEC62-BirA interactome was the presence of ER luminal-resident proteins, including PDIA3, PRDX4, and HSP90B1/GRP94. With the identification of ER luminal proteins limited to the SEC62-BirA reporter line, we initially presumed that the membrane topology of the SEC62-BirA reporter was inverted from the native protein, whose N- and C-termini are cytoplasmic, thereby placing the BirA domain in the ER lumen (Linxweiler et al., 2017; Muller et al., 2010). Alternatively, and given that the ER luminal proteins identified were present in biological triplicates and exceeded significance cutoffs, these data imply that SEC62 is functionally coupled with or proximal to the recently discovered ER luminal protein reflux pathway machinery (Igbaria et al., 2019). To distinguish between these two possibilities, we examined the membrane topology of the SEC62-BirA reporter by protease protection assays, performed on digitonin-permeabilized SEC62-BirA expressing cells (Figure 4E). In this approach, cytosolic domains of ER membrane proteins are expected to be protease accessible, whereas ER lumen proteins are largely protected against protease digestion. GRP94 and TRAP*α*, both ER-resident proteins, were used as proteolysis topology controls. GRP94, an ER luminal protein was protected from proteinase K digestion (Figure 4E). In contrast, TRAP*α* is digested completely at 25 µg/ml of proteinase K (the lowest concentration tested), with detection by a polyclonal antibody raised against the cytosolic domain (Figure 4E). Similar to what we observed with TRAP*α* digestion, anti-BirA reactivity was lost at the lowest proteinase K concentration tested, demonstrating that the SEC62-BirA reporter assumes the membrane topology of the native protein (Figure 4E). To further examine if there was an over-representation of ER luminal proteins in our SEC62-BioID dataset, we assessed the membrane vs. soluble distribution of all 50 interacting proteins using the membranOME database (Lomize et al., 2018; Lomize et al., 2017) (Supplemental Figure 1B). Using this approach, we determined that only 42% of the SEC62 interactome is made up of membrane proteins (Supplemental Figure 1B). This suggests that while we did observe an enrichment of ER luminal proteins, the majority of the unique SEC62 interactors are indeed soluble proteins. Notably, this distribution between membrane vs. soluble protein interactors was mirrored in the set of high-confidence SEC62 interactors identified by the single BioID-reporter experiments, which also include ER luminal proteins (Supplemental Figure 1C, “Type” column). Together, these data further suggest that SEC62 may be proximal to and/or an interactor with an ER luminal protein reflux pathway. Further studies are needed to establish this putative functional link.

### The LRRC59 interactome is enriched in the SRP pathway, ER-resident RNA-binding proteins, and translation factors

Previous *in vitro* studies have shown LRRC59 to interact with the 60S ribosomal subunit, however the biological importance of this interaction and the local LRRC59 membrane environment remains unknown. Following the methodology detailed above, we examined the BioID-LRRC59 interactome. As noted, our analysis identified 25 high-confidence LRRC59 interacting proteins (Figure 5A). Unlike the SEC62-BirA interactome which is enriched for ER functions other than mRNA translation/protein biogenesis, proteins identified in the LRRC59-BirA dataset were highly enriched for functions related to mRNA translation (e.g. eIF2A, eIF5), the SRP pathway (e.g. SRP54, SRP72), and RNA binding (e.g. MTDH, SND1) (Figure 5B-D). Notably, the proteins that had the highest quantitative enrichments for LRRC59-BirA labeling include LRRC59, RRBP1 (p180, ribosome-binding protein), MTDH (AEG-1, an RNA-binding protein), SERBP1 (RNA-binding protein), and SRP72 (SRP protein) (Figure 5C, leftmost heatmap).

**Figure 5.**
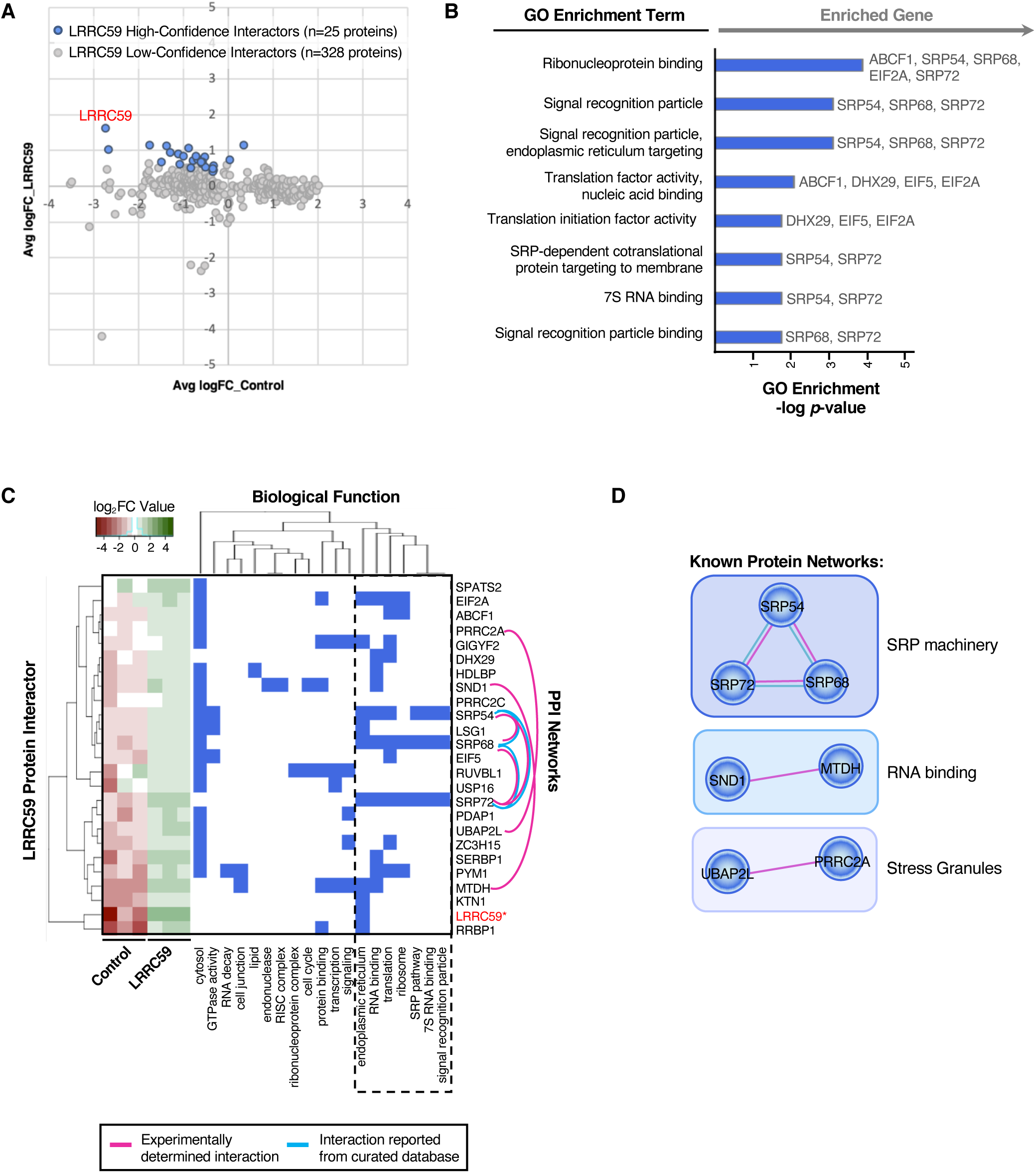
BioID-LRRC59 interacts with SRP pathway, translation machinery, and RNA-binding proteins. **(A)** Comparison of protein abundance for the 353 identified putative interactors in LRRC59-BirA and empty vector control HEK293 cells. Blue dots represent enriched, high-confidence proteins that interact with LRRC59-BirA. Gray dots represent proteins that are less likely *bona fide* interactors of LRRC59-BirA. **(B)** Enriched Gene Ontology (GO) terms associated with high-confidence LRRC59-BirA interactors. Genes assigned to each enriched GO term are listed on the right. **(C)** Clustering of LRRC59 high-confidence interactors based on co-occurrence of functional annotations. The left-most heatmap represents protein abundance values across biological replicates in control and LRRC59-BirA HEK293 cells. Protein-protein interaction (PPI) networks among LRRC59 interacting proteins, as annotated by STRING, are visualized by pink and/or cyan edges. **(D)** Alternative view of PPI networks among LRRC59 high-confidence interacting proteins, based on STRING annotations.

Similar to our previous interactome analyses, we generated a PPI network for the 25 LRRC59-interacting proteins, using STRING (Szklarczyk et al., 2019; Szklarczyk et al., 2017) (Figure 5C-D). This analysis revealed three primary protein networks within the LRRC59 interactome: the RNA-binding proteins MTDH/AEG-1 and SND1 (Sarkar, 2013), the stress granule proteins UBAP2L and PRRC2A (Youn et al., 2018), and importantly the SRP subunit proteins SRP54, SRP68 and SRP72, indicating unusually heavy coverage of SRP (Figure 5C-D, depicted as pink and cyan edges linking interacting proteins). Recently, we reported that MTDH/AEG-1 is an ER-resident integral membrane RBP that predominately binds integral membrane protein-encoding transcripts (Hsu et al., 2018). Importantly, our previous study implicated MTDH/AEG-1 in the localization of secretory and membrane protein-encoding mRNAs to the ER, suggesting that LRRC59 may also bind functionally-related mRNAs. SND1, which has been shown to interact with MTDH during overexpression studies in cancer models, is a tudor domain-containing protein that modulates the transcription, splicing, and stability of mRNAs related to cell proliferation, signaling pathways, and tumorigenesis (Guo et al., 2014; Li et al., 2008). These functional annotations are consistent with models where LRRC59 functions in a complex with MTDH/AEG-1 and SND1 to recruit/regulate mRNAs for translation on the ER membrane. In a similar vein, the BioID data identified LRRC59 as an SRP interactor. SRP is best characterized for its role in the signal sequence-dependent trafficking of ribosomes engaged in the translation of secretory/membrane proteins. Intriguingly, the C-terminus of LRRC59 (located in the ER lumen) shares overlapping sequence structure with the SR receptor (Ohsumi et al., 1993), further implicating LRRC59 function in the SRP pathway and/or translation of secretory/membrane protein mRNAs on the ER. Additionally, the interaction of LRRC59 with the protein-protein network pair, UBAP2L-PRRC2C, may relate to mRNA regulation via stress granule assembly. Stress granules are membrane-less structures formed from non-translating mRNPs during stress (Khong et al., 2017; Protter and Parker, 2016). Stress granules are typically composed of several RNA-binding proteins, along with factors involved in translation initiation and mRNA decay. Interestingly, the LRRC59 interactome is enriched for all three classes of factors. We also report PRRC2C, a known paralog of PRRC2A which is required for the efficient formation of stress granules (Youn et al., 2018), as an LRRC59 interacting protein. These data indicate that stress granule proteins may associate with ER-compartmentalized translation centers (e.g. LRRC59 interactome). Combined with our previous study, these data demonstrate that LRRC59 associates with ER-bound ribosomes and scaffolds a protein interactome highly enriched in SRP pathway machinery and RNA-binding proteins, suggesting a relationship between LRRC59 and stress granule formation. Experiments to test these hypotheses are currently ongoing.

### Orthogonal validation confirms the direct interaction of LRRC59 with mRNA translation-related factors

One limitation to a proximity proteomics approach is that the identified protein interactors cannot be distinguished as stable vs. transient interactors. To determine if LRRC59 stably interacts with SRP machinery, translation factors, and/or RNA-binding proteins, we performed LRRC59 native co-immunoprecipitation (co-IP) studies followed by mass spectrometry. In brief, Caco-2 cells were cultured, detergent extracts prepared, and LRRC59 captured via indirect immunoprecipitation, using an affinity purified anti-LRRC59 antisera. Following mass spectrometric analysis, raw data files were processed with Protein Discoverer and Scaffold to perform semi-quantitative analysis via total spectral counts for the identified proteins. High-confidence interacting proteins of LRRC59 were subsequently identified using CompPASS (Sowa et al., 2009), which is an unbiased, comparative proteomics software platform. In total, 2,678 prey within each IP were identified (Figure 6A), and of these proteins, 102 were determined to be high-confidence interacting proteins (HCIP) of LRRC59 (D-score *≥* 1) (Figure 6B**, Supplemental File S4**). Notably, 20% of these HCIPs overlapped with those determined by BioID (Figure 6B). As expected, these shared LRRC59 targets include SERPB1, DHX29, PRRC2C, SRP68, and LRRC59 itself, which were among the most enriched biotin-labeled proteins within the LRRC59-BirA experiment. We also recovered the other highly enriched proteins SRP72, SRP54, and RRBP1 in the LRRC59 co-IP data (Figure 6E); however, their D-scores (0.95, 0.92, and 0.90, respectively) were just below the conservative threshold. Given that SRP is itself a ribonucleoprotein complex, these data are consistent with SRP acting as a stable member of the LRCC59 interactome.

**Figure 6.**
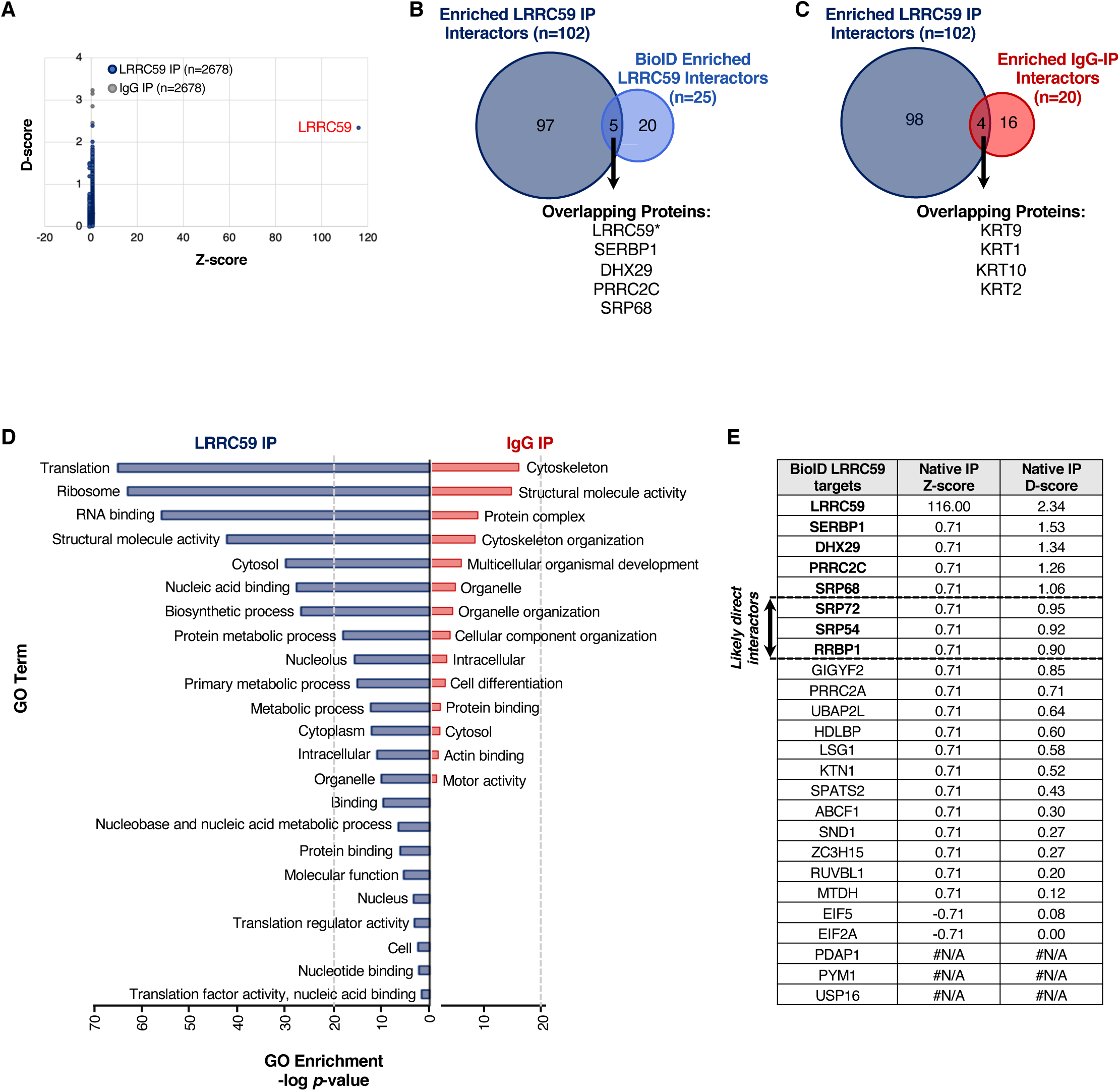
LRRC59 co-IP screen for direct interactions with SRP pathway, translation machinery, and RNA-binding proteins. **(A)** Comparison of D- and Z-scores, as determined by CompPASS analysis, for all proteins identified to interact with LRRC59 (blue) and IgG (control; gray) via immunoprecipitation (IP). Each dot is one of the 2,678 proteins identified by mass spectrometry. **(B)** Number and overlap of enriched, high-confidence interactors of LRRC59, as determined by co-IP (dark blue) or isobaric tagging (BioID; light blue) approaches. **(C)** Number and overlap of high-confidence interactors (D-score *≥* 1) of LRRC59 (dark blue) or IgG (red), as determined by co-IP. **(D)** Enriched Gene Ontology (GO) terms associated with high-confidence LRRC59 interactors (dark blue, left) or IgG interactors (red, right). **(E)** Comparison of D- and Z-scores for each of the 25 LRRC59-interacting proteins, as determined by the BioID approach.

Since co-IP assays are generally accompanied by high background, we also co-immunoprecipitated non-specific IgG as a control. Importantly, analysis of our IgG-IP yielded only 20 HCIPs, which is a small fraction (19%) compared to the LRRC59 interactome (Figure 6C). We did observe an enrichment of keratin proteins as HCIPs in both LRRC59 and IgG interactomes (Figure 6C), and attribute this to common environmental contamination, as has been previously reported (Mellacheruvu et al., 2013). Thus, our data suggests that the primary *bona fide* interactors of LRRC59 are uniquely enriched by co-IP.

To assess the biological functions of all the HCIPs that directly interact with LRRC59 (n=102), we performed GO analysis. Consistent with our observations of the BioID-LRRC59 interactome, HCIPs determined by co-IP were also strongly enriched for proteins with functions related to mRNA translation and RNA binding (Figure 6D, left side). In contrast, non-immune IgG high-confidence interactors were only mildly enriched in common background proteins (Mellacheruvu et al., 2013) (Figure 6D, right side). Therefore, our data collectively demonstrates that LRRC59 directly interacts with SRP machinery, translation initiation factors, and RNA-binding proteins.

## Discussion

While the protein machinery involved in secretory and membrane biogenesis on the ER is well established, it remains unclear how mRNA translation on the ER, including translation of cytosolic protein-encoding mRNAs, is spatially organized in cells. Moreover, our understanding of how the resident ER proteome contributes to mRNA localization, anchoring, and translational control is lacking. In this communication, we examine these questions by characterizing the protein interactomes of known and candidate ER-resident ribosome receptors in the mammalian cell line HEK293. Of particular interest, our data place LRRC59 in a functional nexus for secretory and membrane protein synthesis via interactions with SRP, translation initiation factors, and RNA-binding proteins. Combined, the results of this study reveal new modes of compartmentalized mRNA translation and expand upon the canonical understanding of the SRP pathway.

### Functional domain organization of the ER membrane

The endoplasmic reticulum is a structurally complex organelle known to serve multiple functions, including mRNA translation, protein translocation, protein folding, post-translational protein modifications, lipid biosynthesis, and calcium transport (English and Voeltz, 2013; Schwarz and Blower, 2016). In addition, the ER contains specialized domains dedicated to interactions with other membrane organelles, such as the mitochondria and endosomes, and was recently demonstrated to participate in stress granule and processing body dynamics (Cohen et al., 2018; Lee et al., 2020; Murley and Nunnari, 2016; Wu et al., 2018). With the regulation of dynamic ER morphology and organelle-organelle interactions under active investigation, insights into the spatial organization of the ER membrane and how this higher order is necessary to accommodate its wide-range of biological functions can be expected to provide molecular intersections between the two processes.

Using an unbiased, multiplexed proteomics approach to examine the protein neighborhoods of membrane-bound ribosomes, we identified over 200 proteins in the HEK293 reporter model, many of which clustered into discrete functional categories. Importantly, each of the four tested ER ribosome interactor-BioID reporters had unique sets of interacting proteins, which is consistent with proteins being enriched in functional domains of the ER membrane. In agreement with published structural data, our proximity proteomics study revealed SEC61β to interact with other members of the SEC61 translocon, including SEC61*α* and SEC63. Remarkably, we also discovered SEC61β to interact with 143 (n=145 proteins, total) other proteins, making it the largest interactome identified by our study. Despite its large size, gene ontology analysis of the SEC61β interactome yielded a strong enrichment for membrane and transport proteins, which parallels SEC61β’s primary role in secretory/membrane protein biogenesis. While our list of SEC61β-BirA protein interactors have functions that converge on those expected of the translocon, our analysis also provides new candidate interacting proteins that may function alongside SEC61β; and by extension, suggests alternative mechanisms for mRNA translation via the SEC61 translocon.

Similarly, we characterized the protein interactome of the ER-resident protein, RPN1, which is a subunit of the OST complex and an accessory component of the translocon (Harada et al., 2009; Kreibich et al., 1978b; Nilsson and and von Heinje, 1993; Wild et al., 2018). In contrast to SEC61β, the high-confidence RPN1 interactome was limited, comprising 13 proteins, making it the smallest interactome identified by our study. Nonetheless, we found RPN1 to interact with SSR1/TRAP*α* and SSR3/TRAP*γ*, which has been previously structurally validated (Nilsson et al., 2003; Pfeffer et al., 2014). Members of the OST complex, as well as 60S ribosomal proteins, were also among the list of RPN1 interactors, which is consistent with the spatial assignment of RPN1 and its function in N-linked glycosylation and ribosome binding, respectively. While we did not pursue the direct functional relationship between these proteins and RPN1, our data provides a new platform for studying dynamic regulation of mRNA translation by the OST complex.

### The SEC62 interactome is functionally diverse

The ER-localized members of the SEC gene family have been extensively studied via genetic and biochemical approaches, revealing how Sec61p, Sec62p and Sec63p interact with one another and operate collectively to support translocation of membrane and secretory proteins (Deshaies et al., 1991; Lang et al., 2012; Linxweiler et al., 2017). While these studies have advanced our understanding of the SEC61 translocon and the biological functions of SEC62 and SEC63 in protein translocation, how these proteins interact with the translation machinery, particularly in mammalian cells, has only recently gained attention (Jadhav et al., 2015; Muller et al., 2010). Our system identified SEC62 to interact with 50 proteins. Unexpectedly, the SEC62 interactome was enriched for ER luminal proteins, including BiP, GRP94, PDI, and PRDX4. Identifications of these interactions by both TMT-multiplexed and single reporter proteomics analyses confirmed that the SEC62-BirA reporter has the appropriate orientation at the ER membrane. Moreover, noting that these interactions were not identified in the three other BioID reporters examined, we conclude that these are likely *bona fide* interactions. The existing literature on cytoplasmic and nuclear localizations of ER luminal chaperones such as calreticulin and BiP (Afshar et al., 2005; Duriez et al., 2008; Halperin et al., 2014; Shaffer et al., 2005), along with the recent identification of an ER lumen protein reflux pathway (Igbaria et al., 2019), provide key evidence for a retrograde trafficking pathway for ER luminal proteins across the ER membrane and suggest that SEC62 may functionally intersect with such processes. Further study is needed, however, to understand the molecular basis for the observed SEC62-ER luminal protein interactions.

Surprisingly, the SEC62 interactome also includes proteins functioning in cell-cell adhesion, vesicle transport, signaling pathways, and cytoskeleton formation, indicating that SEC62 may have functions independent of protein biogenesis. For example, SEC62 may be important for ER tubule organization and protein transport to the Golgi apparatus. Interestingly, our data also suggests that SEC62 may have a critical role in multiple signaling pathways. To date, the best characterized ER signaling pathway is the Unfolded Protein Response (UPR). In the UPR, the accumulation of misfolded proteins at the ER triggers a signaling cascade that includes transcriptional (e.g. ATF6) upregulation of ER chaperones (e.g. BiP, protein disulfide isomerases (PDI), GRP94), and ERAD components – which are all represented in our list of interactors. We also found SEC62 to interact with proteins that function in the Wnt and Notch signaling pathways, which are less commonly studied in the context of ER regulation, though it has been reported that Wnt signaling proteins are retained in the ER due to inefficient secretion (Burrus and McMahon, 1995; Moti et al., 2019). To this point, the ER-resident glycoprotein, Oto, regulates Wnt activity by binding Wnt1 and Wnt3a to facilitate its retention in the ER (Zoltewicz et al., 2009). Whether SEC62 acts as another ER-resident protein that binds Wnt-related factors to regulate the accumulation and burst of Wnt ligands remains to be determined. Similarly, our data suggests that SEC62 may play a role in the glycosylation process of Notch proteins, thereby influencing Notch activation. Understanding how SEC62 may function in the UPR, Notch, and/or Wnt signaling pathways has the potential to shed new light on how defects in these signaling cascades at the ER contributes to genetic human disorders.

### A role for LRRC59 in the spatial organization of protein synthesis on the ER

Despite the discovery of LRRC59 decades ago, little is known about its biological function (Hoffman et al., 2019; Ichimura, 1992; Ichimura et al., 1993; Ohsumi et al., 1993; Tatematsu et al., 2015; Xian et al., 2020). Early sequence analysis revealed that the cytoplasmic domain of LRRC59 contains a number of intriguing structural features, including leucine-rich repeats (LRR), which are known protein-protein interaction motifs, hydrophilic regions (KRE), and several regions of charged residues that could serve as sites for protein-protein interactions and ribosome binding activity (Ichimura, 1992; Ohsumi et al., 1993). Indeed, via proximity proteomics, we identified high-confidence interactions with 25 proteins. Importantly, these interactions are likely occurring on the cytosolic domain of LRRC59, as predicted, since the reporter construct places the BirA terminal to the LRR, KRE, and transmembrane-spanning domains (Supplemental Figure 1A). Prominent in the LRRC59 interactome were subunits of the SRP (e.g. SRP54, SRP72, SRP68), translation initiation factors (e.g. eIF2A, eIF5, DHX29), and other ER-RBPs (e.g. SERBP1, MTDH). The prevalence of these interactions link LRRC59 to the regulation of secretory and membrane protein synthesis on the ER. In support of this view, we recently demonstrated that LRRC59-BirA constructs robustly label ER-bound ribosomes (Hoffman et al., 2019), and previous *in vitro* studies demonstrated that LRRC59 binds the 60S ribosomal subunit (Ichimura, 1992; Ohsumi et al., 1993).

Here, we propose six possible mechanisms by which LRRC59 may regulate mRNA translation on the ER membrane (Figure 7). First, given the enrichment of SRP subunits in both our BioID and co-IP experiments, LRRC59 may directly interact with the SRP receptor (Figure 7A) and/or SRP (Figure 7B) to recruit mRNA/ribosome/nascent peptide complexes to the ER membrane for continued mRNA translation. Alternatively, LRRC59 may bind translationally active ribosomes via protein-protein interactions occurring on its large LRR- and KRE-containing cytoplasmic domain (Figure 7C). Given the enrichment of translation initiation factors interacting with LRRC59, our data also suggests that LRRC59 may bind both 60S and 40S ribosomal subunits, as well as initiation factors in proximity to facilitate mRNA translation initiation on the ER membrane (Figure 7D). Another possible mechanism for a LRRC59 function in mRNA translation is via directly binding mRNA (Hsu et al., 2018) and/or indirectly targeting mRNAs through interactions with ER-localized RBPs (Figure 7E). By anchoring localized mRNAs either directly or indirectly, LRRC59 may then recruit ribosomes (as previously postulated) for subsequent mRNA translation (Figure 7E). Finally, our data also reveal LRRC59 to interact with proteins that associate with stress granules (e.g. UBAP2L, PRRC2A, PRRC2C). Therefore, we hypothesize that stress granules may reside proximal to LRRC59 as a mechanism to spatially and temporally fine-tune protein synthesis upon changes in cellular homeostasis (Figure 7F). Although these enriched interactions are highly suggestive of a role for LRRC59 in mRNA translation regulation on the ER membrane, further studies are necessary to provide mechanistic support for these hypotheses.

**Figure 7.**
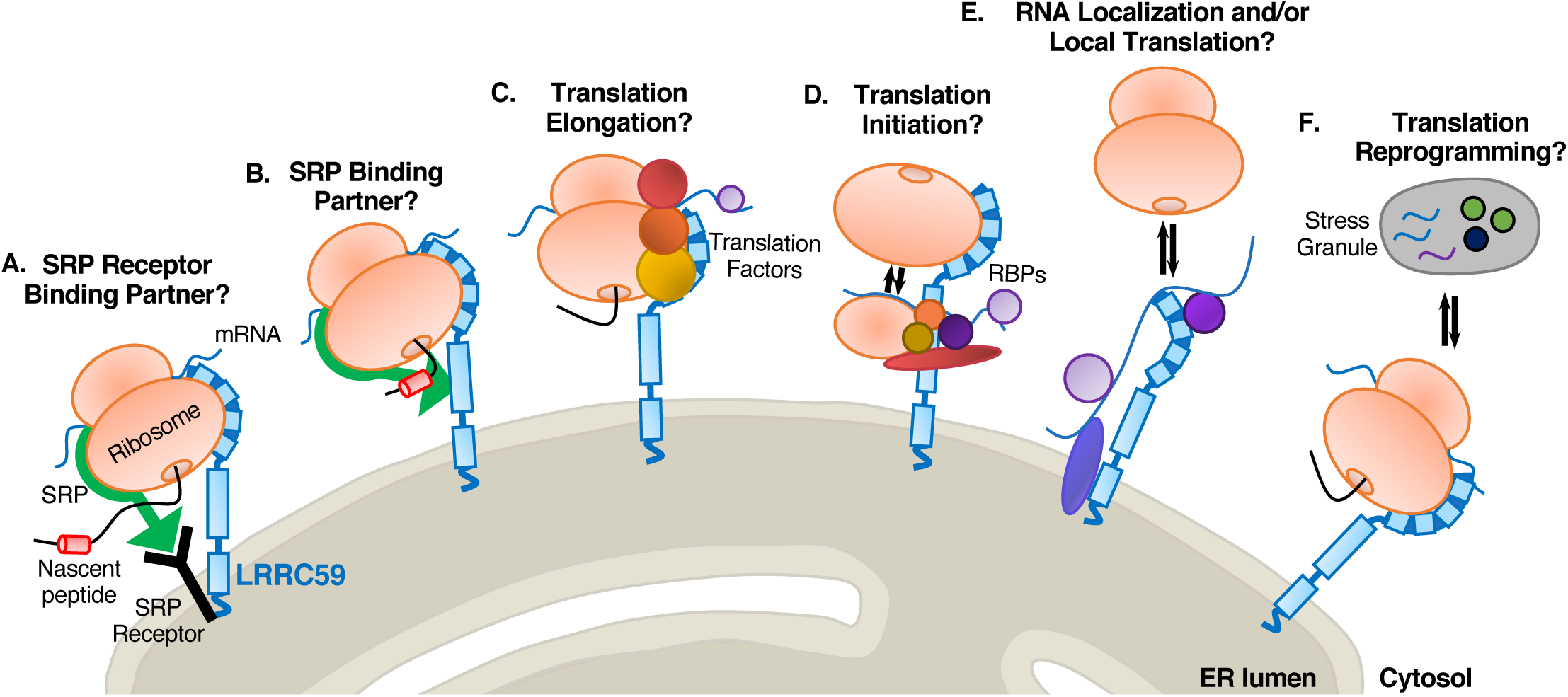
Model depicting LRRC59 interactions with ER localized mRNA translation. Proximity proteomics revealed LRRC59 to significantly interact with SRP factors, translation machinery (including the ribosome), RNA-binding proteins (RBPs), and proteins associated with stress granules. As depicted, LRRC59 may interact with the **(A)** SRP receptor or **(B)** SRP to recruit translationally-engaged ribosomes to the ER membrane for continued mRNA translation. **(C)** LRRC59 recruits mRNA/ribosome/nascent peptide complexes by directly interacting with associated translation factors and/or RBPs, independent of the SRP pathway. **(D)** LRRC59 interacts with the 40S and 60S ribosomal subunits, along with translation factors, RBPs, and mRNA to facilitate translation initiation. **(E)** LRRC59 may anchor mRNAs on the ER membrane via direct RNA binding activity and/or through interactions with other mRNA-bound RBPs, thereby recruiting nearby ribosomes for subsequent mRNA translation. **(F)** LRRC59 may interact with stress granules to fine-tune the activity of translating ribosomes in response to alterations in cellular homeostasis. The depicted modes of mRNA regulation by LRRC59 are not mutually exclusive.

In summary, we demonstrate that ER-resident proteins proximal to bound ribosomes are organized via protein network interactions. We provide evidence that SEC61β interacts with proteins important for secretory and membrane translocation; demonstrate that RPN1 interacts with OST complex subunits and ribosomal proteins; and propose a new function for LRRC59 in regulating mRNA translation of secretory/membrane-encoding proteins via the SRP pathway. These data also reveal an array of possible biological functions that SEC62 may facilitate, including signaling pathways, redox homeostasis, and protein folding. Together, these data offer significant insights into mechanisms of translational organization on the ER and advance understanding into the diversity of functions performed by this central organelle.

## Materials and Methods

### Generation of BioID chimera and Flp-In™ T-Rex™ HEK293 cell lines

BirA-chimera constructs are described in (Hoffman et al., 2019). HEK293 Flp-In**™** T-REx**™** cell lines were generated according to the manufacturer’s instructions (ThermoFisher Scientific). BirA-containing plasmids (0.4 µg), along with the pOGG4 (4 µg) plasmid were transfected into cells using 7.5 µL of Lipofectamine 3000 (ThermoFisher, L3000001). All transfections were performed in 6-well culture dishes at 80% confluency. Colonies were selected for between 48 hours and two weeks post-transfection using 100 μg/mL hygromycin (MediaTech, 30-240-CR, Manassas, VA) and 15 μg/mL blasticidin (ThermoFisher, R21001). A negative control cell line (“Empty Vector Control”) was generated by recombination of an empty vector pcDNA5-FRT/TO and antibiotic selection for an empty vector matched control.

### Sequential detergent fractionation and cell lysis

Cells were washed twice with ice-cold PBS containing 50 μg/mL of cycloheximide (CHX) (VWR, 94271, Radnor, PA) for 3 minutes. To extract the cytosolic (C) fraction, cells were permeabilized for 5 minutes at 4°C in buffer containing 110 mM KOAc, 25 mM HEPES pH 7.2, 2.5 mM MgCl_2_, 0.03% digitonin (Calbiochem, 3004010), 1 mM DTT, 50 μg/mL CHX, 40 U/mL RNAseOUT (Invitrogen, 10777-019, Carlsbad, CA), and protease inhibitor complex (PIC) (Sigma Aldrich, P8340). Supernatants were collected as the cytosolic fraction, and cells were then rinsed with wash buffer (110 mM KOAc, 25 mM HEPES pH 7.2, 2.5 mM Mg(OAc)_2_, 0.004% digitonin, 1 mM DTT, 50 μg/mL CHX, 40 U/mL RNAseOUT, and PIC). To extract the membrane (M) fraction, the washed cells were then lysed for 5 minutes at 4°C in 400 mM KOAc, 25 mM HEPES pH 7.2, 15 mM MgCl_2_, 1% NP-40, 0.5% DOC, 1 mM DTT, 50 μg/mL CHX, 40 U/mL RNAseOUT, and PIC. Subcellular fractions were cleared by centrifugation (15,300 x *g* for 10 minutes). Total cell lysis was performed by incubating cells at 4°C for 10 minutes in membrane lysis buffer (as listed above), followed by centrifugation at 15,300 x *g* for 10 minutes.

### BirA labeling of microsomes

Canine pancreas rough microsomes (RM) (Walter and Blobel, 1980) were adjusted to a concentration of 4 mg/mL in 500 μL of BirA reaction buffer (20 mM Tris pH 8, 5 mM CaCl_2_, 100 mM KCl_2_, 10 mM MgCl_2_, 3 mM ATP, 1.5 mM biotin, 5 mM phosphocreatine (Sigma-Aldrich, P7936-1G), and 5 μg/mL of creatine kinase (Sigma-Aldrich, C3755-3.5KU)). Purified recombinant BirA*-GST fusion protein was added at a concentration of 10 μg/mL. Following 0, 1, 3, 6, and 18 hours, 100 μL of reaction was removed, flash frozen in a dry ice/ethanol bath, and stored at –80°C for subsequent analysis.

### Immunoblotting

Protein lysate concentrations were determined using a Pierce BCA Protein Assay Kit (ThermoFisher, 23225). Proteins were resolved by SDS-PAGE in 12% acrylamide gels containing 0.5% of trichloroethanol. Gels were UV irradiated for 5 minutes and imaged using an Amersham Imager 600 (GE Life Sciences). Gels were then equilibrated in Tris-glycine transfer buffer for 5 minutes and transferred onto nitrocellulose membranes. Membranes were blocked in 3% BSA and probed for BirA (Abcam, ab14002), Streptavidin-RD680 (Li-Cor, P/N 925-68079; 1:20,000), TRAPα (Migliaccio et al., 1992), or GRP94 (Jagannathan et al., 2011). Membranes were incubated with isotype-matched secondary antibodies (Li-Cor, Lincoln, NE; 1:10,000), and imaged by infrared fluorescence detection using the Odyssey Clx (Li-Cor), where signal intensities were quantified by densitometry analyses. To examine total protein levels, immunoblots were stained with either India Ink or Ponceau S solution (Sigma-Aldrich).

### Protease protection assay

The SEC62-BirA construct was expressed overnight as reported in (Hoffman et al., 2019). Cultures were then placed on ice, permeabilized in digitonin-supplemented cytosolic buffer (as described above), rinsed, and incubated with cytosolic buffer containing 0, 25, or 50 µg/mL of Proteinase K (Bioline) for 30 minutes at 4°C. Protease digestions were quenched by addition of 0.5 mM PMSF. Cell extracts were prepared and immunoblots performed as above.

### TMT/Isobaric tag mass spectrometry

#### Sample preparation and proteolytic digestion

Three biological replicates from each reporter cell line were affinity isolated on streptavidin magnetic beads, eluted in 120 µL of biotin elution buffer (2% SDS, 20 mM biotin, 2 M thiourea, 0.5 M Tris unbuffered), and prepared according to the standard S-Trap digestion protocol (Protifi, Inc.; (Yang et al., 2018)). Briefly, each sample was loaded onto its respective S-Trap column, washed four times with S-trap binding buffer (90% MeOH, 100 mM TEAB), and digested by adding 0.8 µg of sequencing grade trypsin to the top of each S-trap tip with incubation for one hour at 47°C. The peptides were eluted from the S-trap tip first with 50 mM TEAB, then with 0.2% aqueous formic acid, and finally with 50% acetonitrile in 0.2% aqueous formic acid. The peptide elutions were combined and dried via SpeedVac. Peptide yield from each sample was determined to be approximately 20 µg based on BCA Protein Assay (ThermoFisher Scientific).

#### TMT labeling

Dried samples, determined to contain approximately 22 µg of digested peptide each, were brought to room temperature and resuspended in 70 µL 200 mM TEAB. An aliquot (20 µL) from each of the 15 samples was combined to make a Study Pool QC (SPQC). TMT reagents (TMT10Plex plus TMT11-131C, Product A37725) were dissolved in 45 µL acetonitrile for 5 minutes with vortexing. Labeling reagent (20 µl) was added to each sample for 2 hours at room temperature. Sample labeling was then quenched with 4 µL of 5% v/v hydroxylamine in 200 mM TEAB for 15 minutes. The TMT samples for each set were combined into a 1.5 mL Eppendorf tube, acidified to 1.0% formic acid, frozen, and lyophilized to dryness overnight.

#### Pre-fractionation

Each TMT labeled peptide set was fractionated to improve depth of proteome coverage using a Pierce High pH Reversed-Phase Peptide Fractionation Kit (ThermoFisher Scientific, Part 84868). The fractionation was performed according to the manufacturer’s protocol and yielded 8 peptide fractions for analysis. Water/acetonitrile mixtures with 0.1% v/v triethylamine (TEA), pH 10, were used for reversed-phase fractionation. 5% v/v wash was used to remove excess TMT reagent, then fractions were collected at 10, 12.5, 15, 17.5, 20, 22.25, 25, and 50% v/v MeCN. These fractions were independently acidified to 1% formic acid and dried via SpeedVac. Samples were subsequently resuspended in 22 µL 1/2/97 v/v/v TFA/MeCN/water.

#### Liquid chromatography – tandem mass spectrometry

Approximately 1 µg of TMT-labeled peptide from each fraction was analyzed by nanoscale liquid chromatography – tandem mass spectrometry (LC-MS/MS) on a nanoAquity UPLC (Waters) coupled to an Orbitrap Fusion Lumos Tribrid mass spectrometer (ThermoFisher Scientific). Peptides were first trapped on a column at 99.9% water and 5 µL/min, followed by separation at 0.4 µL/min on an analytical column (Waters Corporation) with a gradient from 3 to 30% MeCN (0.1% formic acid) over 90 minutes. Column eluent was introduced to the MS via electrospray ionization (+2.1kV) and a source temperature of 275°C. Upon easy-IC internal mass calibration, tandem MS sequencing and quantification was performed using a full-scan spectrum at 120k resolution, followed by MS2 sequencing at 50k resolution with HCD fragmentation at 38 V. MS/MS was performed with an isolation width of 0.7 Da, a cycle time of 1 second until the next full scan spectrum, and 60 seconds dynamic exclusion. Raw data and *.mgf peaklist files for this study have been uploaded to the MASSive data repository and are available at (https://massive.ucsd.edu/).

#### TMT-labeled MS data processing

Raw MS data (ftp://massive.ucsd.edu/MSV000085009/) was converted to *.mgf format using Proteome Discoverer v2.1 (ThermoFisher Scientific) and submitted to Mascot v2.5 (Matrix Sciences, Inc.) for database searching. Peptide matching included 5 ppm precursor and 0.02 Da product ion tolerance, fixed carbamidomethyl (C), along with variable modifications TMT10 (N-term, K) and deamidation (N, Q). Searches were performed against the curated human proteome (www.uniprot.org), plus common contaminant sequences such as ALBU_BOVIN, ADH1_YEAST, ENO1_YEAST, and BIRA_ECOLI. A reverse-sequence decoy database was appended for False Discovery Rate (FDR) determination. Scaffold Q+ v4.8.5 (Proteome Software, Inc.) was used to quantify TMT-label based peptide and protein identifications. Peptide identifications were accepted if they could be established at greater than 50.0% probability by the Scaffold Local FDR algorithm, while protein identifications were accepted if they could be established at greater than 99.9% probability and contained at least 1 identified peptide. Protein probabilities were assigned by the Protein Prophet algorithm (Nesvizhskii et al., 2003). TMT reporter ion channels were corrected based on isotopic purity in all samples according to the algorithm described in i-Tracker (Shadforth et al., 2005). Normalization was performed iteratively (across samples and spectra) on intensities, as described in (Oberg et al., 2008). Spectra data were log-transformed, pruned of those matched to multiple proteins and those missing a reference value, and weighted by an adaptive intensity weighting algorithm. Relative protein abundance across the experiment was expressed as the log_2_ ratio to the reference (SPQC) channel average for all samples (**Supplemental File S1**). Percent missing values were calculated at the protein level for the SPQC channels, as well as all channels. A *p*-value using a Student’s t-test was then calculated comparing each biological group (n=3) versus the SPQC (n=6).

#### Identification of interaction networks

A combination of statistical prioritization, 2D clustering, and principal components analysis (PCA) was used to identify putative interaction networks from the dataset. The data were curated such that proteins only quantified in one TMT set, or missing in more than 40 of the total channels were excluded from consideration (86 of 1,263 proteins). A *p*-value was then calculated using a Student’s t-test between each BirA-fusion sample group (n=3) and the SPQC group (n=6) to determine whether a protein was statistically different in each biological group (BirA reporter) from the average of all groups (SPQC). Proteins that did not pass a Bonferroni-corrected *p*-value < 0.1 (raw *p*-value < 1e-4) were removed, yielding 353 proteins as putative interactors **(Supplemental File S2)**. Finally, putative interaction networks were identified using unbiased 2D hierarchical clustering (Robust, Ward’s Method) in JMP 14.0 (SAS Institute, Cary NC). The clustering analysis only included the BirA interactome samples (not the SPQC samples) in order to reduce the potential for cluster mis-assignment.

### Label-free proteomic analysis of BioID proteomes

#### Sample preparation

For single BioID reporter studies, reporter cell culturing, reporter expression, cell fractionation, detergent lysis, and affinity isolation of biotinylated proteins was performed as above. Samples were subjected to one dimensional SDS-PAGE. 25 µL of sample was combined with 5 µL of 100 mM DTT and 10 µL of NuPAGE™ (ThermoFisher Scientific) 4X loading buffer, and samples were then heated to 70°C for ten minutes with shaking. SDS-PAGE separation was performed using 1.5 mm 4-12% Bis-Tris pre-cast polyacrylamide gels (Novex, ThermoFisher Scientific) and 1X MES SDS NuPAGE™ Running Buffer (ThermoFisher Scientific), including NuPAGE™ antioxidant. SDS-PAGE separation was performed at a constant 200V for five minutes, gels fixed for 10 minutes, stained for 3 hours, and destained overnight following manufacturer instructions.

#### Gel band isolation and trypsin digestion

Gel bands of interest were isolated using a sterile scalpel, transferred to protein LoBind tubes (Eppendorf), and minced. Gel pieces were washed with 500 µL of 40% LCMS grade acetonitrile (MeCN, ThermoFisher Scientific) in AmBic, with shaking at 30°C. Gel pieces were shrunk with LCMS grade MeCN, the solution discarded, and the gel pieces dried at 50°C for 3 min. Reduction of disulfides was performed using 100 µL of 10 mM DTT at 80°C for 30 min with shaking, followed by alkylation with 100 µL of 55 mM IAM at room temperature for 20 min. This liquid was aspirated from the samples and gel pieces were washed twice with 500 µL AmBic. LCMS grade MeCN was added to shrink the gel pieces in each sample, then samples were swelled in AmBic, and this process was repeated. The gel pieces were shrunk a final time by adding 200 µL of LCMS grade MeCN, and heating for 3 min at 50°C to promote evaporation. Trypsin digestion was performed by addition of 30 µL of 10 ng/µL sequencing grade trypsin (Promega, Madison, WI) in AmBic followed by 30 µL of additional AmBic. The samples were incubated overnight at 37°C with shaking at 750 rpm. Following overnight digestion, 60 µL of 1/2/97 v/v/v TFA/MeCN/water was added to each sample and incubated for 30 min at room temperature and 750 rpm to extract peptides, and the combined supernatant was transferred to an autosampler vial (Waters). Gel pieces were shrunk in 50 µL additional MeCN for 10 min to extract the maximum number of peptides, and combined with the previous supernatant. The samples were dried in the Vacufuge (Eppendorf) and stored at −80°C.

#### Qualitative analysis of gel electrophoresis samples

All gel band samples were resuspended in 20 µL of 1/2/97 v/v/v TFA/MeCN/water and analyzed by nanoLC-MS/MS using a Waters nanoAcquity LC interfaced to a Thermo Q-Exactive Plus via a nanoelectrospray ionization source. 1 µL of each gel band sample was injected for analysis. Each sample was first trapped on a Symmetry C18, 300 μm x 180 mm trapping column (5 µL/min at 99.9/0.1 v/v H2O/MeCN for 5 minutes), after which the analytical separation was performed using a 1.7 μm ACQUITY HSS T3 C18 75 μm x 250 mm column (Waters). The peptides were eluted using a gradient of 5-40% MeCN with 0.1% formic acid at a flow rate of 400 nL/min with a column temperature of 55°C for 90 minutes. Data collection on the Q Exactive Plus mass spectrometer was performed with data dependent acquisition (DDA) MS/MS, using a 70,000 resolution precursor ion (MS1) scan followed by MS/MS (MS2) of the top 10 most abundant ions at 17,500 resolution. MS1 was performed using an automatic gain control target of 1e6 ions and maximum ion injection (max IT) time of 60 ms. MS2 used AGC target of 5e4 ions, 60 ms max IT time, 2.0 m/z isolation window, 27 V normalized collision energy, and 20 s dynamic exclusion.

#### Single reporter MS data processing

Database searching was performed as described by TMT-labeled MS data processing. For single reporters, data was searched using trypsin enzyme cleavage rules and a maximum of 4 missed cleavages, fixed modification carbamidomethylated cysteine, variable modifications biotinylated lysine, deamidated asparagine and glutamic acid, and oxidized methionine. The peptide mass tolerance was set to +/− 5 ppm and the fragment mass tolerance was set to +/− 0.02 Da. False discovery rate control for peptide and protein identifications was performed using Scaffold v4 (Proteome Software, Inc.) **(Supplemental File S3)**.

### Native LRRC59 immunoprecipitation and mass spectrometry

#### Sample preparation

Caco-2 cells were cultured according to ATCC recommendations and processed at ca. 90% confluence. Cell extracts were prepared by addition of 0.5 mL per 15 cm plate of NP-40 lysis buffer (1% NP-40, 100 mM KOAc, 50 mM HEPES pH 7.2, 2 mM Mg(OAc)_2_, PIC, 1 mM DTT). Lysates were maintained on ice for 20 minutes and cleared by centrifugation (10,000 x *g*, 10 minutes). The supernatant fractions were diluted 1:1 in dilution buffer (50 mM HEPES, 100 mM KOAc, 2 mM Mg(OAc)_2_, PIC, 1 mM DTT) and supplemented with 5 µg/mL of LRRC59 antibody (A305-076A, Bethyl Labs, Montgomery TX) or rabbit IgG (Sigma-Aldrich, St. Louis, MO). Samples were incubated with end-over-end rotation overnight at 4°C. Dynabead Protein G beads (ThermoFisher, Waltham MA) were added to a concentration of 30 µL/mL and rotated for 2 hours at 4°C. Beads were washed 3x in buffer 1 (0.1% NP-40, 100 mM KOAc, 50 mM HEPES pH 7.2, 2 mM Mg(OAc)_2_, PIC, 1 mM DTT), 1x in buffer 2 (0.1% NP-40, 500 mM KOAc, 50 mM HEPES pH 7.2, 2 mM Mg(OAc)_2_, PIC, 1 mM DTT), and 1x in PBS. Proteins were eluted in an equi-bead volume of 2x Laemmli buffer by heating at 70°C for 20 minutes and submitted for mass spectrometry analysis.

#### LRRC59-IP data analysis

Raw MS data (.sf3 files) were processed using Scaffold 4 Proteome Software (Proteome Software, Inc.) to obtain total spectral counts for each sample. Protein interactors of LRRC59 and IgG (control) were then determined by performing *CompPASS* (Sowa et al., 2009), which is an unbiased, comparative proteomics analysis. Any prey in each IP with a D-score greater than or equal to one was considered to be a high-confidence interacting protein **(Supplemental File S4)**.

### Bioinformatic analyses

#### Gene Ontology

GO analyses were performed using the Cytoscape application, BiNGO (Maere et al., 2005), with a Benjamini and Hochberg FDR correction (significance level of 0.05) to enrich for terms after multiple testing correction. A custom set of genes expressed in our multiplexed BioID experiment was used as background for examination of SEC62-BirA and LRRC59-BirA interactors, while the entire human annotation (provided within the application) was used as a reference background for LRRC59 interactors determined by native IP. Additional functional information (as depicted by the heatmaps/matrices and protein color-coding) was extracted by batch querying each set of protein interactors against the MGI (Bult et al., 2019; Krupke et al., 2017; Smith et al., 2019) and STRING (Szklarczyk et al., 2019; Szklarczyk et al., 2017) databases.

#### Protein-Protein Interaction Networks

Protein-protein interaction analyses of SEC62-BirA (n=50) and LRRC59-BirA (n=25) interactors were performed using the STRING database (Szklarczyk et al., 2019; Szklarczyk et al., 2017). Only experimentally determined interactions and those reported from a curated database were considered.

#### Identification of membrane proteins

The list of SEC62-BirA interactors (n=50) was intersected with a membrane protein annotation file downloaded from the MembranOME database (Lomize et al., 2018; Lomize et al., 2017). Of the 50 SEC62-interacting proteins, 21 (42%) were identified as membrane proteins. Membrane protein classification was validated by manually searching each of the 50 proteins against The Human Protein Atlas (Uhlen et al., 2015; Uhlen et al., 2010).

## Acknowledgements

We thank the Duke University School of Medicine Proteomics and Metabolomics Shared Resource for their invaluable proteomics services, along with members of the Nicchitta laboratory, in particular Jessica Childs, Jason Arne, and JohnCarlo Kristofich, for their helpful feedback on this project. This work was supported by NIH grant GM101533 **t**o C.V.N. The authors declare no competing financial interests.

## Author Contributions

Conceptualization and experimental design: A.M. Hoffman and C.V. Nicchitta; Investigation, formal analysis, and visualization: M.M. Hannigan, A.M. Hoffman, C.V. Nicchitta, J.W. Thompson, T. Zheng; Writing – original draft: M.M. Hannigan and C.V. Nicchitta; Writing – review and editing: A.M. Hoffman, J.W. Thompson; Funding acquisition, project administration, and supervision: C.V. Nicchitta,

## Figure Legends

**Supplemental File. Results from Mass Spectrometry Analyses.** Excel file with results of **(S1)** identified proteins (n=1263) and **(S2)** putative protein interactors (n=353) from the isobaric/tandem mass spectrometry analyses of the multiplexed BioID reporter constructs; **(S3)** protein interactors determined by single, label-free BioID reporter mass spectrometry analyses; **(S4)** and high-confidence protein interactors (n=102) of LRRC59 as determined by mass spectrometry analyses of native LRRC59-immunoprecipitations.

---

**Supplemental Figure 1.**
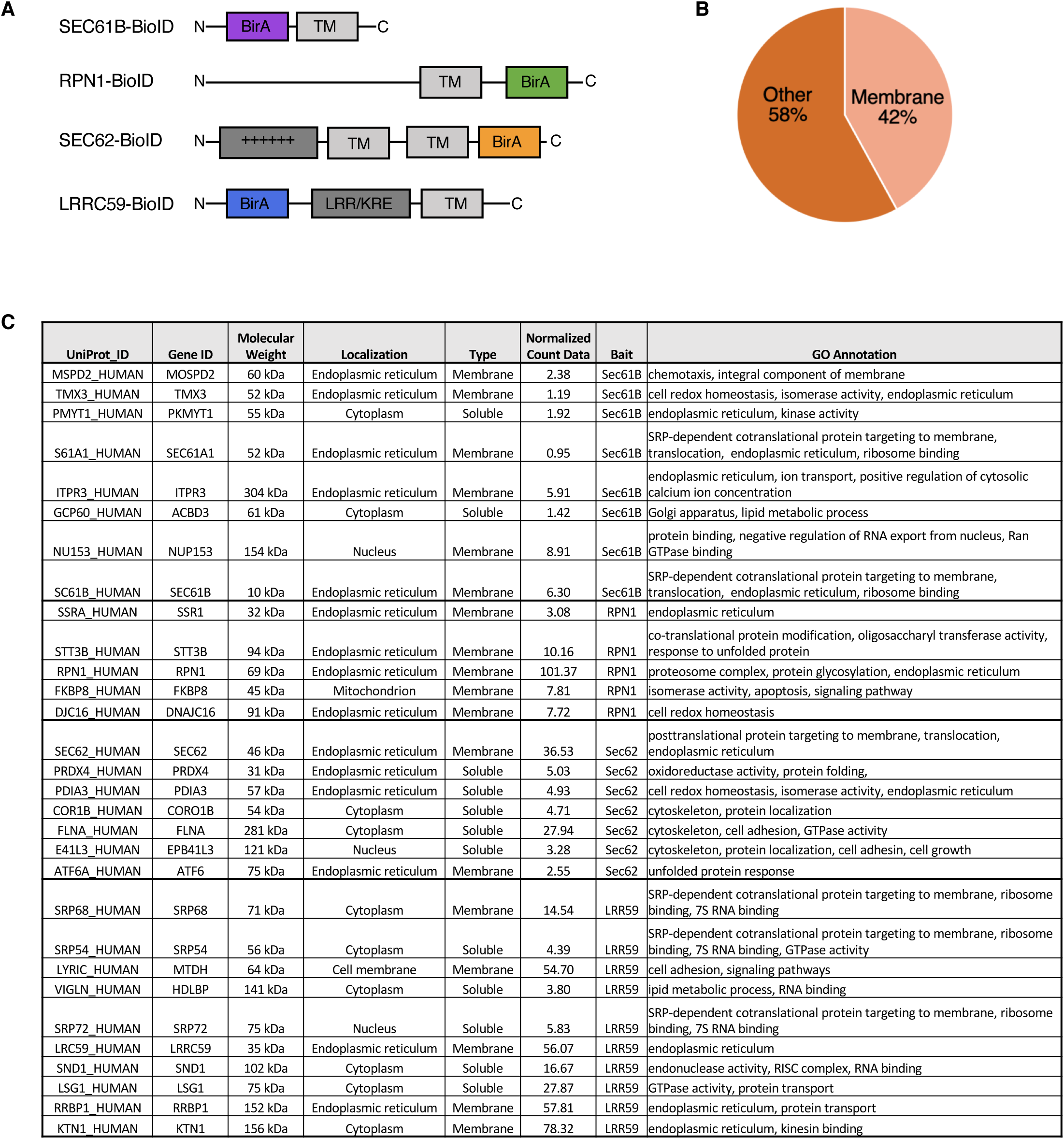
Classification of BioID-chimeras and their associated interactomes. **(A)** Schematic of BirA-containing reporter constructs. **(B)** Distribution of membrane proteins identified within the SEC62-BirA interactome, based on membranOME annotations. **(C)** Subset of enriched, high-confidence interactors of BioID-SEC61β, -RPN1, -SEC62, or -LRRC59, as determined by single reporter, label-free mass spectrometry analyses.

